# BoardION: real-time monitoring of Oxford Nanopore Technologies devices

**DOI:** 10.1101/2020.06.09.142273

**Authors:** Aimeric Bruno, Jean-Marc Aury, Stefan Engelen

## Abstract

**Summary:** One of the main advantages of the Oxford Nanopore Technology (ONT) is the possibility of sequencing in real time. However, the ONT sequencing interface is not sufficient to explore the quality of sequencing data in depth and existing tools do not take full advantage of real-time data streaming. We present BoardION, an interactive web application, dedicated to sequencing platforms, for real-time monitoring of all ONT sequencing devices (MinION, Mk1C, GridION and PromethION). BoardION allows users to easily explore sequencing metrics and optimize the quantity and the quality of the generated data during the experiment. It also enables the comparison of several flowcells to evaluate the library preparation protocols or the quality of input samples.

**Availability:** Source code and a Docker image are available at http://www.genoscope.cns.fr/boardion/

## 1 Introduction

Since 2014, Oxford Nanopore Technologies (ONT) has launched a range of nanopore sequencing devices (MinION, Mk1C, GridION and PromethION). The ONT technology allows the sequencing of long DNA molecules with a read accuracy of currently 96% (Tyler et al. 2018) and already has a wide range of applications (Jain et al. 2018, Belser et al. 2018, Simpson et al. 2017, Sessegolo et al. 2019). Unlike traditional next-generation sequencing platforms, which provide data at the end of the sequencing, ONT sequencing data is generated in real time. Users can access time-critical information (Petersen et al. 2019) and thus be able to wash and reuse the flowcell once the result is obtained (Euskirchen et al. 2017). However, the ONT interface (MinKnow) does not provide enough plots or interactivity to explore the quality of sequencing data in depth. In addition, existing quality control tools such as MinIONQC (Lanfear et al. 2019), pycoQC (Leger et al. 2019) and NanoPack (De Coster et al. 2018) do not take full advantage of real-time data sequencing as they analyze the sequencing data after the run ends. Here, we introduce BoardION, an interactive web application for real-time monitoring of ONT sequencing runs. BoardION offers the possibility for sequencing platforms to remotely and simultaneously monitor all their ONT devices. It also allows the comparison of several flowcells and assess the library’s preparations or the quality of the input samples.

## 2 Software description

BoardION is organized in two components, a preprocessing script and a web application. The first component uses the information contained in the sequencing summary file generated by the guppy basecaller. Since this file can be quite large, a C++ preprocessing program treats the new lines at each execution to generate lightweight datasets with the metrics of interest. Thanks to this preprocessing, statistics can be computed and updated in the web application every minute. The second component is a web application accessible from anywhere regardless of the user’s configuration. The application, developed in R, uses ggplot2 (Wickham 2009), R shiny (Chang 2019) and R plotly (Carson 2018) to create dynamic JavaScript visualization. Documentation, github and dockerhub repositories, as well as an interactive demo version of BoardION are available at http://www.genoscope.cns.fr/boardion/.

## 3 Real-time monitoring

The BoardION application computes metrics such as yield, translocation speed, read length and quality. These metrics are displayed through dynamic and customizable plots cumulatively from the start of the sequencing run or every ten minutes to monitor the experiment in real time. We add a channel-view of the flowcell to assess the presence of bubbles or contaminants, which can reduce the run efficiency. During the experiment, the consumption of reagents leads to a decrease in the translocation speed, read quality and yield (Tyler et al. 2018). ONT recommends keeping the speed above 300 bases per second (ONT 2019). BoardION’s « Run in progress » tab (Fig 1A) allows us to monitor the sequencing metrics in real-time and add reagents (refueling) at the right time to avoid an irreversible drop in efficiency of a PromethION run (Fig 1 E, F, G, H). Thanks to the BoardION monitoring, the run generates twice as much data (81 Gb instead of 43 Gb). Monitoring experiments with BoardION allows sequencing platforms to set metric thresholds below which the flowcell must be refueled or washed to sequence a new sample.

**Fig. 1.**
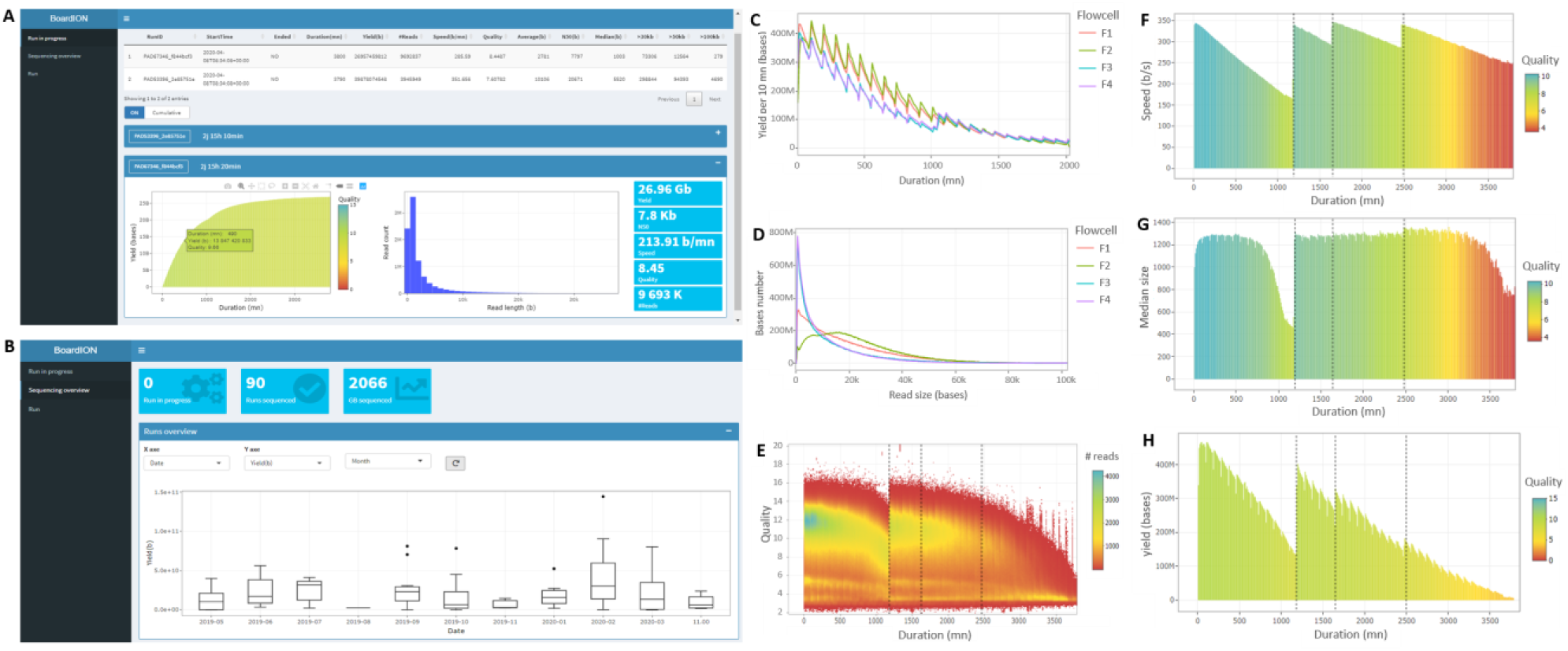
Examples of BoardION tabs (A and B) and plots (C - H). Plots C and D illustrate a comparison of two strains of the same species sequenced with different library sizes. (C) shows that the yield is not the same between the two strains (green and red lines compared to purple and blue). Library preparation like green flowcell is the best for generating long reads (D). Plots E, F, G and H illustrate real-time monitoring of a refueled promethION flowcell. The variation of the chosen metrics is represented as a function of the elapsed time (every ten minutes). Plot H shows the number of reads (color) generated at a given time (x) with a particular quality (y). We have added dotted black lines highlighting the refueling time (1200, 1650 and 2450 mn). The plots show a drop of the read median size (G) and quality (E) after 500 mn of sequencing at a speed behind 260 b/s (F). The first refuel at 1200 mn restores initial speed (F) and median size (G) but also an almost initial yield (400Mb versus 460Mb, plot H). The last two refuels were made at a speed of around 300 b/s to avoid a drop in sequencing metrics.

The « Sequencing overview » tab of the BoardION application offers the possibility of comparing the results of several flowcells. The first panel gives an overview of the metrics for all the sequencing runs (Fig 1B) and allows users to follow the evolution of the sequencing efficiency according to new releases (protocols, flowcells, sequencers or basecallers). Next panels are flexible and users can interactively choose flowcells and metrics and add them to current plots. These panels follow the evolution of a given metrics and flowcells over time. This functionality is particularly useful to compare different preparation protocols or sequencing efficiency of different strains of the same species (Fig 1C, 1D).

## 4 Conclusion

BoardION is a web application for real-time monitoring of ONT sequencing runs. The application can be installed via the Docker distribution. BoardION’s dynamic and interactive interface allows users to explore sequencing metrics easily and quickly and to optimize in real time the quantity and the quality of the generated data. It also offers the possibility of comparing several sequencing experiments which can be useful to follow the evolution of nanopore technology. We believe it will be useful to the community in helping sequencing platforms establish the best sequencing guidelines for different types of samples.

